# Heat flows solubilize apatite to boost phosphate availability for prebiotic chemistry

**DOI:** 10.1101/2024.08.06.606769

**Authors:** Thomas Matreux, Almuth Schmid, Mechthild Rappold, Daniel Weller, Ayşe Zeynep Çalışkanoğlu, Kelsey R. Moore, Tanja Bosak, Donald B. Dingwell, Konstantin Karaghiosoff, François Guyot, Bettina Scheu, Dieter Braun, Christof B. Mast

## Abstract

Phosphorus is an essential building block of the most prominent biomolecules, such as polynucleic acids, and has likely played that role since the beginning of life. Despite this importance for prebiotic chemistry, phosphate could not be supplied by the atmosphere, and had to be fueled mainly by geological phosphate sources. However, phosphorus was scarce in Earth’s rock record and often bound in poorly soluble minerals, with the calcium phosphate mineral apatite as key example. While specific chemical boundary conditions that bind calcium have been used to address this so-called phosphate problem, a fundamental process that solubilizes and enriches phosphate from geological sources remains elusive. Here, we show that ubiquitous heat flows through rock cracks can liberate phosphate from apatite by the selective removal of calcium. Phosphate’s surprisingly strong thermophoresis not only achieves its 100-fold up-concentration in aqueous solution, in particular it also boosts its solubility by two orders of magnitude. We show that the heat-flow-solubilized phosphate can feed the synthesis of trimetaphosphate, increasing the conversion 260-fold compared to the thermal equilibrium case. Heat flows thus enhance solubility as a geological parameter to unlock apatites as phosphate source for prebiotic chemistry, providing a key element in solving early life’s phosphate problem.

---

Phosphorus is an integral part of all life and is used in metabolites such as nucleoside-polyphosphates, phospholipids in cell membranes, and the backbone of DNA or RNA^1^ (Fig. 1a). The abundance of phosphorus in biomolecules is likely to be no accident but, on the contrary, dictated by function. Oligonucleotides, for example, become negatively charged under neutral conditions through the phosphate backbone, which is essential to keep them readable and prevent hydrolysis^2^. Given this central importance, it is assumed that phosphate must have been available at an early stage during the origins of life.

**Figure 1.**
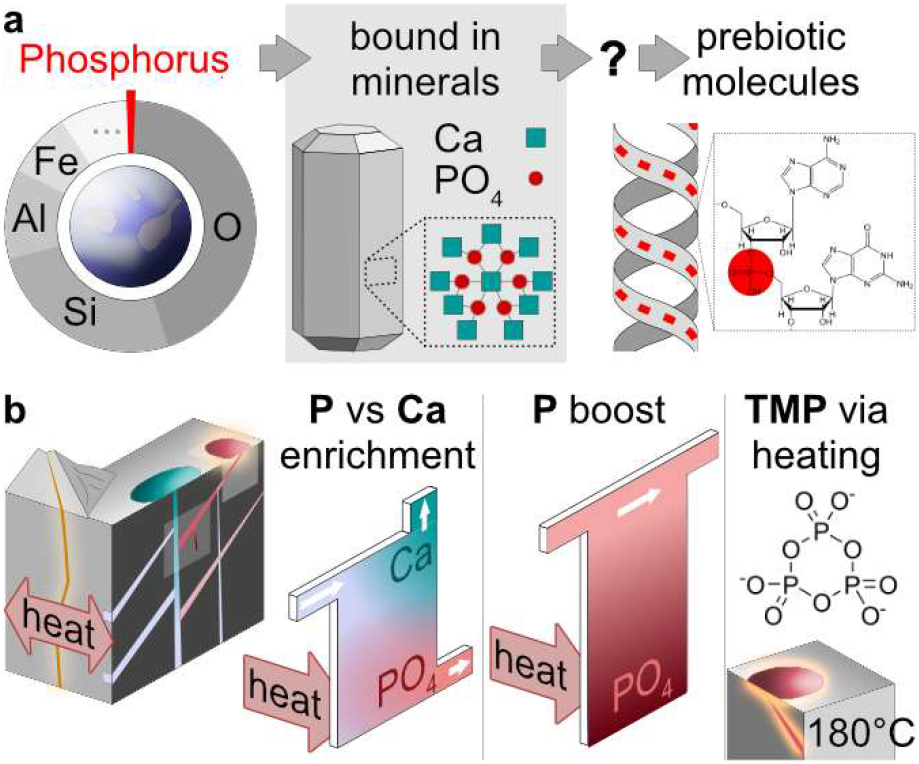
A geothermal solution to the phosphate problem on the early Earth. **(a)** Phosphorus only constitutes around 0.1 wt % of Earth’s crust and is mostly bound as phosphate in apatite minerals, which renders it inaccessible for nascent life. **(b)** Heat flows in geothermal systems are able to enrich phosphate against calcium, boosting phosphate solubility at neutral pH and its absolute concentrations for downstream synthesis of energy-rich trimetaphosphate (TMP).

However, in the presence of divalent ions^3^, phosphate is water-insoluble over a wide pH range and, most importantly, precipitates with calcium to form apatite Ca_5_(PO_4_)_3_(F,Cl,OH) or brushite CaHPO_4_·2 H_2_O. These processes likely reduced the amount of free and reactive phosphate 4 billion years ago, keeping it far below the high millimolar concentration regime required for prebiotic chemistry^4–8^. This high demand also stems from the poor reactivity of orthophosphate, e.g., for the phosphorylation of nucleosides^4–6^ and precursors^7,8^, efficient reaction buffering^4,8^, and the synthesis of reactive phosphate species^8–11^. In combination with its low availability, this constitutes the so-called phosphate problem, which has been identified as one of the major hurdles to understanding the origins of life on Earth^2,12^.

Apatite was presumably relatively abundant compared to other phosphate-bearing minerals^12–16^, for instance in frequent inclusions in igneous, sedimentary, and metamorphic rocks^15,16^. Therefore, strategies to release its bound phosphate for use in prebiotic chemistry have been investigated. While apatite easily dissolves under low pH conditions (1.5 – 2.5), as found in lacustrine environments, acidic lakes^17,18^, hot springs^19^, and hydrothermal systems^20–23^ with steep pH gradients^24–27^, such environments are often incompatible with prebiotic chemistry, which commonly requires neutral to alkaline conditions^17,23,27^. Here, the dissolved phosphate quickly precipitates with presumably abundant calcium, so that despite the broad range of pH conditions available in geological systems, one is back at square one of the phosphate problem.

Chemically, this dilemma has been addressed experimentally with the help of chelating agents such as ammonium oxalate^28^ or citric acid^29^, which bind calcium and thus keep phosphate in solution even under neutral to alkaline conditions. At the same time, it raises the question of the plausibility of the high chelating agent concentrations required^30^. In carbonate lakes, sequestration of free calcium into calcium carbonate is possible, with acidic inflows constantly supplying new phosphate and evaporation increasing its concentration to 0.1 molal^31^. However, evaporation unselectively concentrates all ions, which is potentially detrimental to prebiotic chemistry reactions^32–34^ and fatty acid vesicle^32^ or coacervate stability^35^.

The low reactivity of phosphate motivated the search for more reactive and soluble phosphorus-containing molecules. Reduced phosphorus of meteoritic origin has been shown to phosphorylate glycerol^36^ and yield diamidophosphate^37^. Condensed phosphates such as cyclic trimetaphosphate (TMP)^38–41^ have been shown to trigger peptide synthesis^40,42,43^ or the phosphorylation of nucleosides^40,44^. Chemically activated phosphates such as phosphoenolpyruvate^8^ and acetyl^10^, carbamoyl^11^, and imidazole phosphate^9^ have been demonstrated to drive efficient phosphorylation of a variety of prebiotically relevant species, but also require high phosphate concentrations for their synthesis.

Thus it becomes apparent that prebiotic chemistry would benefit massively from a chelate-independent and widely accessible process that drives the large-scale release of orthophosphate from geomaterials, protects it from diffusive dilution and enables its downstream condensation to reactive polyphosphates. A promising scenario to consider here is permeable pathways in rocks, such as water-filled fractures in magmatic or geothermal environments, as well as sedimentary layers of shallow submarine or lacustrine settings that are exposed to heat fluxes. Such thermomicrofluidic environments have been investigated as a simple but ubiquitous and versatile tool that drives molecular selection and prebiotic reactions^43^. Here, the superposition of thermally driven thermophoresis and convection could have enriched prebiotic building blocks depending on their charge, size and type, ultimately boosting downstream prebiotic chemistry through the shift of solute compositions towards reactive species^34,43,45^.

Heat sources are widespread as a universal consequence of the second law of thermodynamics^46^, e.g., found near meteoritic impacts^47^, in volcanic environments^48^, or in hydrothermal systems^49^. This raises the question whether heat flows could offer a new pathway for broad prebiotic availability of phosphate from minerals and other geomaterials.

Here, we show that heat flows through thin rock fractures can selectively shift the composition of solutions from phosphate-rich minerals such as apatite and thus prevent their precipitation under neutral pH conditions. Molecule-selective thermogravitational enrichment, thus, has a similar effect as chemical approaches that actively remove the calcium responsible for phosphate precipitation (Fig. 2). While apatite dissolves in acidic flows, the resulting solution would re-precipitate in shallow, neutral waters under thermal equilibrium conditions. We show that the heat-flow-driven enrichment can deliver 15 millimolar of solubilized phosphate from apatite at neutral pH, even forming highly reactive TMP under mild heating. We extend this concept to a wide range of further geomaterials, such as clays and sands, and show that the highly dilute phosphate in their leachates can be thermophoretically accumulated within porous niches more than 100-fold (Fig. 3). Our experimental findings are consistent with geochemical modeling^50,51^ and demonstrate an active coupling of geological leaching and precipitation effects with thermal non-equilibrium systems, thereby unlocking a widely available source for phosphate on the prebiotic Earth.

**Figure 2.**
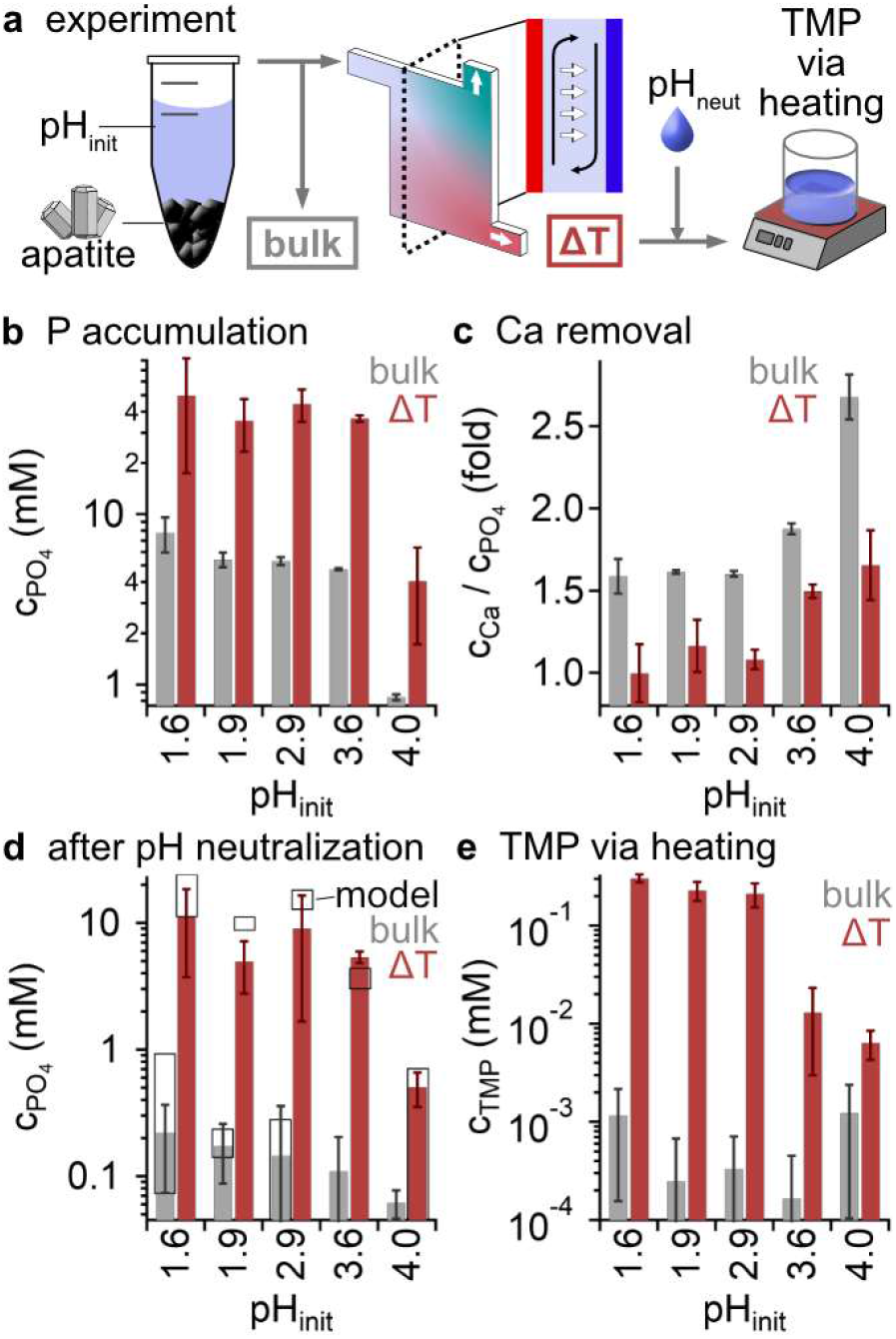
Heat-flow driven solubilization of phosphate from apatite. **(a)** Experiment. Acidic-dissolved phosphate (pH 1.6 to pH 4), is flushed through a heat flow chamber, leading to the selective enrichment of phosphate at the bottom outlet by the interplay of convection (black) and thermophoresis (white). Downstream neutralization mimics the transition to prebiotic chemistry conditions that allow, for instance, the formation of TMP. **(b)** Dissolved phosphate was accumulated from initially low concentrations (bulk, grey) and extracted in the bottom outlet of the heat flow chamber (ΔT, red). **(c)** Here, calcium concentrations were depleted relative to phosphate by the thermal non-equilibrium, which shifts the Ca:PO_4_ ratio from 5:3 found in apatite (bulk, grey) to 1:1 (ΔT, red) (see Supplementary Fig. 2). **(d)** Under the neutral conditions required for prebiotic chemistry, previously acidic-dissolved phosphate and calcium precipitated (see Supplementary Fig. 3-4). In contrast to the bulk case (grey), the heat-flow-driven removal of calcium (ΔT, red), boosted the solubility of phosphate up to 100-fold. The results were verified by geochemical modeling (black boxes, see Methods). **(e)** Mild heating to 180 °C triggered the formation of highly reactive TMP from the heat-flow-altered solutions (ΔT, red), increasing yields more than 100-fold compared to the absence of thermal gradients (bulk, grey).

**Figure 3.**
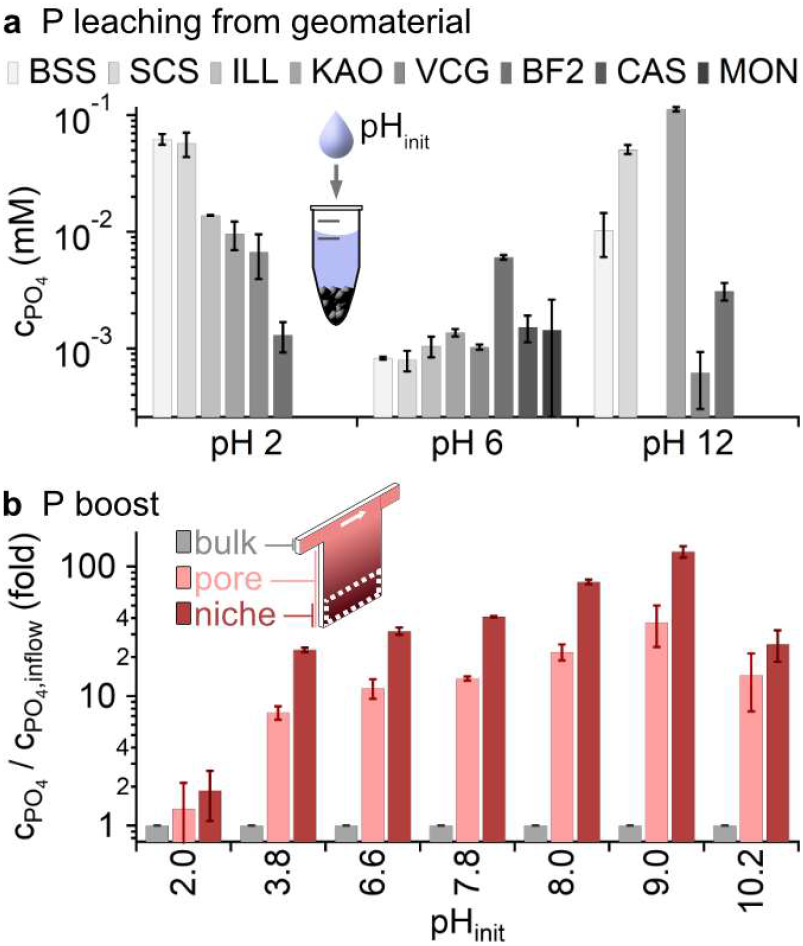
Phosphate-rich habitats formed by heat-flow-driven accumulation. **(a)** Leaching from geomaterials in solutions of different pH_init_ yielded low phosphate concentrations: BSS: basalt sand, SCS: Siliclastic sand, ILL: illite, KAO: kaolinite, VCG: volcanic glass, BF2: basalt F2, CAS: carbonate sand, MON: montmorillonite (for SEM images and compositions see Supplementary Fig. 5-6 and Supplementary Table 3). **(b)** Heat flows accross water-filled fractures boosted the phosphate concentrations from the top-feeding inflows (“bulk”) by a factor of 130-fold at the pore bottom (“niche”), or 40-fold averaged over the whole crack (“pore”).

## Results

Apatite and especially fluorapatite are assumed to be among the most abundant phosphate sources on the early Earth^12–14^. However, the solubility of phosphate from apatite strongly depends on pH, peaking at acidic conditions that are widely incompatible with prebiotic chemistry^4,6–8,10^, and is essentially insensitive to temperature, mass-to-volume ratio, and grain size (Supplementary Fig. 1)^52^. To obtain an experimental baseline without the effect of heat-flow-driven enrichment, we first characterized the amount of phosphate leached from natural apatite under bulk conditions^34,52^. We tested ground apatite samples with different compositions from three locations (Ipirá complex, Brazil, Durango, Mexico and Ontario, Canada, see Supplementary Table 1) and exposed it to a solution with a pre-set pH. After reaching its equilibrium in pH and solute concentrations (Supplementary Fig. 1), we found the resulting phosphate concentrations to be in agreement with previous literature studies^52^.

In contrast to such closed laboratory systems, the geological systems we consider here are assumed to have an open structure and would therefore maintain a constant pH at a leaching-independent, externally-defined value through acidic geothermal flows^17–23^. Although a larger quantity of dissolved phosphate would be expected in such open systems at a fixed calcium-to-phosphate ratio of 5:3, precipitation with calcium under near-surface, the more neutral conditions required for prebiotic chemistry^17,23,27^ would lead to vanishing phosphate concentrations in solution.

To study this scenario experimentally, we prepared 5 samples in which ∼3 g of apatite (using the Brazil apatite, see Supplementary Table 1) was dissolved by repeatedly supplying HCl to reach the respective predefined initial pH values (shown in Fig. 2). This procedure resulted in the apatite-specific calcium-to-phosphate ratios at increased absolute concentrations (Fig. 2B,C grey bars). Given that a certain degree of dilution is expected in an open system due to the geothermal flows on its way to the surface, we have approximated this effect by diluting the above samples 4-fold (see Methods). We mimicked the transition to neutral, near-surface pH conditions by repeated addition of NaOH until a constant neutral pH was reached (see Fig. 2D, Supplementary Fig. 3 and Methods, Eqs. 1-3). As expected, the resulting phosphate concentrations were at mere micromolar levels due to its nearly-complete precipitation with the calcium ions present (Fig. 2D, grey bar). The precipitation equilibria (Fig. 2D, black hollow boxes) were also modeled using PHREEQC-software^50,51^, yielding a good agreement with the experiment findings. Analysis of precipitates by SEM and geochemical modeling^50,51^ revealed the formation of calcium phosphates such as apatite (see Supplementary Table 2, Supplementary Fig. 3-4). Different final pH conditions (8, 10, 12) and temperatures (30 °C, 60 °C, 90 °C) that may occur in a natural setting resulted in the same precipitation characteristics and phosphate concentrations (see Supplementary Fig. 3).

### Heat-flowen-driven solubilization

The characteristics of this flow system massively changed in the presence of heat flows that spatially separate phosphate and calcium ions before reaching the more neutral environments near the surface, where unaltered apatite-leachates would precipitate completely. We experimentally implemented such a setting as may be plausibly expected to be found in heated rock cracks^47–49^ using an open microfluidic heat flow chamber with a thickness of 200 µm across and applied a thermal gradient of 20 K (see Fig. 2A). In this flow-through system, we set the inflow volume rate of acidic-dissolved apatite solution to 15 nl/s^53^ and the outflow volume rate in the two outlets to 0.75 nl/s (bottom) and 14.25 nl/s (top), respectively. As discussed below, more shallow thermal gradients in larger pore networks are expected to lead to the same results.

We explored pH conditions for apatite dissolution between 1.6 and 4 to study the respective characteristics of phosphate enrichment in triplicate experiments and extracted the outflows for post-analysis. As shown before^26,34,43^, moderate flow rates did not disturb the selective accumulation of ions through concurrent solvent convection (black arrows) and solute thermophoresis (white arrows).

These experiments showed that phosphate and calcium were selectively enriched in the respective outlet channels. As depicted in Supplementary Fig. 2, in the bottom outlet, phosphate was accumulated up to 70 % stronger than calcium. Accordingly, the thermal non-equilibrium shifted the phosphate-to-calcium ratio from the apatite-defined 3:5 to 1:1 (see Fig. 2C). This local excess of the anionic species was balanced by a local reduction of pH, providing charge neutrality (Supplementary Fig. 2).

As for the unaltered solution, we mimicked the passage of the heat-flow-enriched phosphate solution to the pH-neutral surface by successively adding NaOH until equilibrium was reached at pH 7-9. The precipitation of phosphate and calcium still occured, far too few calcium ions were present to bind all the dissolved phosphate and thereby remove it from solution. This led to a significantly higher final phosphate concentration of 15 mM compared to the case without heat flow-induced phosphate enrichment with 0.2 mM (Fig. 2D). The results were consistent with geochemical modeling ^50,51^ using measured concentrations of all participating ions obtained by ion chromatography (see Methods).

Thus, heat-flow-driven ion enrichment yielded a 100-fold increase in phosphate solubility, offering new opportunities towards the synthesis of activated phosphate species^8–11,25,38,40,41,44^. To test this, we heated the neutral phosphate solutions obtained before, which were previously thermally fractionated (Fig. 2C “ΔT”) or not altered (Fig. 2C “bulk”), to 180 °C and measured the amount of TMP formed. Efficient synthesis of TMP usually requires high temperatures as found in volcanic environments (>500 °C). Our comparably mild but also prebiotically more widely available heating only resulted in 1 µM TMP concentrations in the solutions not altered by heat flow (Fig. 2D “bulk”). In contrast, up to 0.3 mM TMP formed in the heat-flow-enriched solutions, showing a 260-fold boost in concentrations (Fig. 2D “ΔT”). While without heat-flows, only 0.05 % of the initially acidic dissolved phosphate was converted to TMP due to its re-precipitation with calcium, the thermal non-equilibrium boosted the conversion to up to 12 % of the starting material (see Methods, Eq. 4). As TMP has been shown to accumulate efficiently through thermogravitational accumulation due to its high Soret coefficient^43^, the synthesized TMP could be used downstream for TMP-driven prebiotic reactions such as the in-situ dimerization of Glycine^42,43^.

### Establishment of phosphate-rich geo-habitats

However, nature could not afford to rely purely on pure apatite as the sole source of phosphate, which is why we investigate how heat flows could utilize and concentrate phosphate in geothermal streams fed by a variety of phosphate-bearing, igneous, sedimentary, and metamorphic rocks that contained lower amounts of phosphate^54^.

We, therefore, studied the leaching from various basalts, sands, and clays (SEM images of geomaterials are shown in Supplementary Fig. 5-6 and compositions in Supplementary Table 3). Without thermal gradients, the resulting phosphate concentrations barely exceeded 0.1 mM (Fig. 3A). Leaching from basaltic sand (BSS) showed to be most efficient for acidic pH (pH 2); for neutral pH, basalt F2 (BF2) provided most phosphate, amounting to 60 µM. The low orthophosphate concentrations found were insufficient to drive common prebiotic reactions and so far only known to be enhanced by drying at gas interfaces, which is, however, restricted to near-surface environments. We, therefore, aimed to explore how heat-flow-driven accumulation could boost phosphate concentrations in ubiquitous liquid-only settings.

For this, we explored thermogravitational accumulation in small, protected pockets, that could provide reaction niches for prebiotic chemistry. We used the same experimental setup as above, but closed the bottom outlet, creating a water-filled pore connected to a continuous flow of liquid at the top end. We speculated that the closed pore which lacks a lower material outflow would maximize the heat-flow-driven phosphate accumulation. In the experiment, we applied a volume flow rate of 30 nl/s of dilute phosphate (500 µM, Fig. 3B “bulk”) in solutions of various pH over the course of a week. After this timespan, we froze the entire chamber and segmented the frozen content along the height of the chamber into four fractions of equal volume. IC measurement of all fractions revealed an up to 130-fold concentration increase of orthophosphate in the lowest fraction (Fig. 3B “niche”), with an up to 40-fold boost of phosphate over the entire pore (Fig. 3B “pore”). These results demonstrate that heat flows can boost orthophosphate concentrations in all-liquid environments without the need for dry-wet cycles or gas-water interfaces, making this process a geologically widely accessible method for prebiotic chemistry.

## Discussion

In this work, we have studied the heat-flow-driven solubilization of phosphate from natural apatite and other phosphate-bearing geomaterials and its subsequent up-concentration to form geological, phosphate-rich habitats for prebiotic chemistry. To tackle this crucial question, we have taken into account the wide range of pH conditions found in such geological systems: While apatite could be dissolved without heat flows under acidic conditions, the more neutral pH conditions required for prebiotic chemistry would lead to close-to-complete precipitation, rendering the phosphate inaccessible for downstream reactions. However, once the apatite solution was exposed to heat fluxes before reaching neutral conditions, the selective thermophoretic removal of the co-dissolved calcium lead to a 100-fold increase in phosphate solubility. We demonstrated that the achieved phosphate concentrations of up to 15 mM enhanced the formation of reactive TMP, important for driving various prebiotic reactions. The same enrichment mechanism boosted even the low phosphate concentrations that would be obtained from further phosphate-bearing geomaterials by a factor of 130. Such geomaterials also support the formation of polyphosphates and can even enhance the formation of TMP up to 6-fold on kaolinite (Supplementary Fig. 7), a clay also known to increase the phosphorylation of glycerol^55^.

While the temperature gradients used in the experiments were comparably high (20 K) to reduce experimental timescales^56^, it is known that the effect of heat-flow-driven enrichment scales with the system size, ultimately balancing lower thermal gradients in a natural system^34,43^. In addition, steep local temperature gradients can occur in narrow cracks close to wider fractures that host rapid geothermal flows which act as a continuous heat source^53^. Simple heat flows in geothermal environments thus create a protected niche where the established high phosphate concentrations from solubilized apatite and other geomaterials could drive reactions^4–8^, act as a buffer^4,8^, and yield activated phosphate species^8–11^.

## Acknowledgments

The authors thank P. Aikkila for experimental support and P. Aikkila, A. Dass and J. Langlais for fruitful discussions. This work was funded by the Deutsche Forschungsgemeinschaft (DFG, German Research Foundation) under Project-ID 364653263 – CRC 235 (T.M., A.S., D.W., D.B.D., B.S., D.B., C.B.M.), under Project-ID 521256690 – CRC 392 (T.M., B.S., D.B., C.B.M.), under Project-ID 201269156 – SFB 1032 (M.R., D.B.) and under Germany’s Excellence Strategy EXC-2094-390783311 (T.M., A.S., D.B., C.B.M.). Funding from the Volkswagen Initiative ‘Life? – A Fresh Scientific Approach to the Basic Principles of Life’ (T.M., A.Z.C., D.B.D., D.B., C.B.M.) is gratefully acknowledged. The work is supported by the Center for Nanoscience Munich (CeNS).

## Author contributions

Conceptualization: TM, AS, CBM; Methodology: TM, AS, DB, CBM; Investigation: TM, AS, MR, DW, AZC, KRM, KK, CBM; Visualization: TM, CBM; Funding acquisition: DBD, BS, DB, CBM; Supervision: CBM; Writing – original draft: TM, CBM; Writing – review & editing: TM, AS, MR, DW, AZC, KRM, TB, DBD, KK, FG, BS, DB, CBM

## Data, code and materials availability

All data are available in the main text or the supplementary materials. The code used to simulate precipitation dynamics is supplied in Supplementary Data 1.

## Methods

### Materials

NaH_2_PO_4_, Na_3_P_3_O_9_, Na_5_P_3_O_10_, and CaCO_3_ were purchased from Sigma Aldrich (USA), MSA (Methanesulfonic acid), Na_4_P_2_O_7_, NH_4_Cl, NaCl, MgCl_2_, and CaCl_2_ from CarlRoth GmbH (Germany). For all experiments, ion chromatography water was used (Fisher Scientific, USA). For calibration and reference, Dionex Seven Anion Standard II, and Dionex Combined Six Cation Standard II from Thermo Fisher (USA) were used.

### Apatite sample preparation

The composition of apatite samples is shown in Supplementary Table 1. The samples were crushed with a hammer to an average grain size of 1–2 cm and ground by a vibration mill to <500 µm mesh size. The different fractions were separated by hand sieving to grain sizes between 500 and 63 µm. The different apatite samples were characterized using XRF and SEM/EDX (for details, see below), as depicted in Supplementary Table 1-2. The H_2_O_2_ washed samples used in Supplementary Fig. 1 were incubated in H_2_O_2_ (10 vol%) for seven days, rinsed with purified water, and dried for 24 h at 200 °C in a furnace (similar to^34^).

### Major and trace element analyses of apatite samples

For the apatite samples, major and trace element analyses were carried out at University of Mainz (Germany) using a Malvern PANalytical Axios Fast X-ray fluorescence spectrometer (XRF) (Spectris Plc, UK). The major element analyses were carried out on fused glass discs, the trace element analyses were carried out on compacted powder pellets. Typical accuracy of the analyses of the standard references was approximately 1% relative (RMS) for major elements and 4% relative (RMS) for trace elements.

### Preparation of acidic apatite solution

Large stocks of Brazil apatite were mixed with IC grade water and adjusted multiple times with HCl to the desired pH. We waited for 2-3 weeks between adjustments until the pH was steady. Before the experiments, solutions were diluted with three parts of IC grade water (adjusted to experimental pH) and filtered (0.22 µm).

### pH measurement

The pH values were measured using a Thermo Scientific™ Orion™ 8220BNWP pH Electrode (Thermo Fisher Scientific, USA).

### Heat flow cell construction and setup

200 μm thick FEP film (Holscot, The Netherlands) was cut into the designed microfluidic shape with an industrial plotter (CE6000-40 Plus, Graphtec, Germany) and sandwiched between two sapphires (Kyburz, Switzerland) of thickness 500 μm (cooled sapphire, with four laser-cut holes of 1 mm diameter) and 2000 μm (heated sapphire, without holes). The sapphire-FEP-sapphire sandwich was then placed on an aluminum base, with an additional layer of heat-conductive graphite foil (EYGS091203DP, 25 μm, 1600 W/mK, Panasonic, Japan) between the aluminum base and the sandwich, and fixed there with a steel frame and torque-controlled screws for homogeneous force distribution. A second heat-conductive graphite foil (EYGS0811ZLGH, 200μm, 400 W/mK, Panasonic, Japan) ensured the thermal connection between the heated sapphire and the electrical heating element, which was again connected with torque-controlled steel screws. The thickness of the microfluidic chamber was then measured using a confocal micrometer (CL-3000 series with CL-P015, Keyence, Japan). Next, the chamber was pre-flushed with low-viscosity fluorinated oil (3M™ Novec™ 7500 Engineered Fluid) to check for tightness and to drive out gas inclusions. The assembled chamber was mounted (with an intermediate layer of 200 µm heat conducting graphite foil, see above) onto an aluminum block which itself was connected to a cryostat (Grant R5 and TXF200, Grant Industries, UK) for cooling. The heaters were connected to a 400 W 24 V power supply and solid-state relays controlled by Arduino boards running the open-source Repetier firmware. For details on optimizing the individual elements and construction, see also^34,43^.

### Heat flow cell experiments (general)

The inputs and outputs of the heat flow cell were connected via tubings to syringes placed on high-precision syringe pumps (neMESYS 290N low-pressure syringe pump with low-pressure quad syringe holder, Cetoni, Germany). The following microfluidic connections were used (all Techlab, Germany): Connectors (Connector inch, UP P-702-01), End Caps (Tefzel cap for 1/4-28 nut, UP P-755), Screws (Nut, Delrin, flangeless, VBM 100. 823-100.828), Ferrules (Ferrule VBM 100.632) and Tubing (Tubing Teflon (FEP), KAP 100.969). The chemically resistant syringes used were acquired from Göhler HPLC syringes, Germany: 2606714, 2606814, 2606914, 2606015, 2606035, 2606055, and 2606075 (ILS, Germany). The cryostat was set to -30 °C and the heating elements to 95 °C, resulting in temperatures of 20 °C and 50 °C on the cold and hot sides of the sapphire microfluidic chamber, respectively and translating to a temperature difference of 20 K between the inner surfaces of the sapphires^34^. Temperatures were measured on the respective outer surfaces of the sapphires using a thermal imaging camera (ShotPRO thermal imaging camera, Seek Thermal, USA). Before starting the experiments, all tubes and the thermal chamber were rinsed with fluorinated oil, and samples were loaded into the inlet tubes.

### Heat flow cell experiments (separation, Fig. 2)

For separation, an inflow of 15 nl/s was applied, and the flow rates of the syringe pumps (controlled via neMESYS UserInterface, cetoni, Germany) were selected such that 5 % of the inflow was taken from the lower outlet and 95 % from the upper outlet. The experiments were run for three days. Stopping experiments was achieved by setting applied temperatures to room temperature. Samples were then removed from the tubings and collected for ion chromatographic measurement and further experiments.

### Heat flow cell experiments (up-concentration, Fig. 3)

In order to start experiments, the lower outlet was closed, and an inflow of 30 nl/s was applied with an applied temperature gradient as described above. The experiments were run for one week. The experiments were stopped by setting the applied temperatures to room temperature and disconnecting the tubings. The chamber was then frozen at -80 °C for at least one hour. By opening the frozen chamber and sequentially melting 25 % fractions from bottom to top (also see ^43^), local concentrations were measured by ion chromatography. The pore concentration (Fig. 2) is the average of all four fractions.

### Ion chromatography

Samples were injected using an autosampler (AS-DV, ThermoFisher Scientific, USA) and simultaneously measured in two ion chromatography systems.

Measurement of cations was done using an ion chromatography system (Dionex Aquion, ThermoFisher Scientific, USA) with an analytical column (Dionex IonPac CS12A), guard column (Dionex IonPac CG18) and suppressor (Dionex CDRS 600). The chromatography method was set to provide 0.25 ml/min flow using isocratic elution with 20 mM MSA, 15 mA suppression, a cell temperature of 40 °C, and a column temperature of 35 °C. Eluted ions were detected with a conductivity detector (DS6 Heated Conductivity Cell).

Measurement of anions was done using a separate ion chromatography system (Dionex Integrion, ThermoFisher Scientific, USA) with an analytical column (Dionex IonPac AS16 2mm), guard column (Dionex IonPac AG16 2mm), suppressor (Dionex ADRS 600 2mm), eluent generator (EGC 500 KOH) and trap column (Dionex CR-ATC 600). Here, the method comprised a gradient elution starting with 57.5 mM KOH (for 10 min), a linear increase to 62.5 mM KOH over 2 min, isocratic elution with 62.5 mM KOH for 5 min, a direct step to 57.5 mM, and equilibration for 8 min. The flow was set to 0.25 ml/min, with the suppression current set to 47 mA. The cell temperature was set at 40 °C and the column temperature at 35 °C. Eluted anions were measured with a conductivity detector (DS6 Heated Conductivity Cell).

Data was analyzed using Chromeleon 7.2.10 (ThermoFisher Scientific, USA). Calibration was done using standard solutions.

### Re-neutralization of solution

For each re-neutralization step towards a target pH *pH*_*target*_, the pH of the solution *pH*_*sol*_ was measured, and the following concentration of NaOH was added:

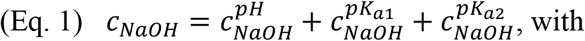

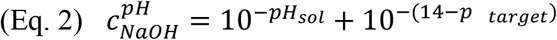

as compensation of the pH difference and

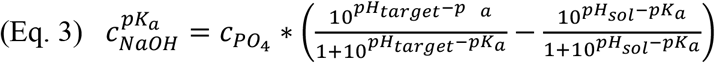

to overcome the pK_a_s of phosphoric acid at 2.2 (*pK*_*a1*_) and 7.2 (*pK*_*a2*_). Phosphate concentrations were measured using ion chromatography (see above).

We incubated all samples after pH adjustment for two weeks at 60 °C to reach the precipitation equilibrium. As this process leads to a shift in the pH, we repeated this procedure until no further precipitation occurred at a static pH. Three neutralization steps proved sufficient in the experiments, validated by modeling (Fig. 2D, Supplementary Fig. 3).

### Geochemical modelling

PHREEQC^50^ was used for modelling, both in its standalone version (3.7.3) and in phreeqpython from Vitens^51^. The code is supplied in Supplementary Data 1.

### SEM-EDX analysis of precipitates and of geomaterials

Particles were carbon coated and measured with a Hitachi SU-5000 SEM (Japan). Samples were imaged in Back-Scattered Electron (BSE) mode in a high vacuum and elements were measured by Energy Dispersive X-ray spectroscopy (EDX). Acceleration voltage was set to 15 and 20 kV, Working Distance (WD) ∼10 mm, at 240000 nA emission current. Element mappings were performed over selected areas with additional point measurements to distinguish between individual crystals.

### Synthesis experiments of trimetaphosphate

Experiments were done in glass vials (17374073, ThermoFisher, USA) which were filled with 10 µl of sample and heated to 180 °C in an oven (Memmert UNB 100 Oven, Germany). After three days, the vials were retrieved, cooled to room temperature, eluted in 500 µl ion chromatography water, and measured by ion chromatography. In the experiments shown in Supplementary Fig. 7, 30 mg of geomaterial was added to the vials before heating with an otherwise identical protocol.

### Other geomaterial samples

BF2: Basalt F2 sample from Kilauea volcano in Hawaii. ILL: Illite. KAO: Kaolinite clay, purchased from Sigma Aldrich (USA). MON: Montmorillonite clay, purchased from Sigma-Aldrich (USA). ZEO: Zeolite, obtained from Zeocem (Slovakia). CAS: Carbonate sand, a mixture of carbonate ooids and other grains that were collected in the Bahamas, containing fragments of foraminifera and other carbonate shells. BSS: Basalt sand, crushed up, iron-rich basalt. SCS: Siliclastic sand, quartz-rich beach sand containing iron minerals purchased from Sigma Aldrich (USA). VCG: Volcanic glass, crushed and powered obsidian. SEM images are shown in Supplementary Fig. 5-6 and compositions obtained by EDX analysis are described above in Supplementary Table 3.

### Conversion of trimetaphosphate (TMP) from initial acidic-dissolved phosphate

To obtain the conversion to TMP of the initially acidic-dissolved phosphate, we calculated:

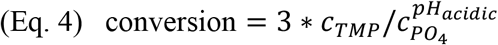

(as each TMP contains three phosphates).

### Leaching experiments (Supplementary Fig. 1)

Samples were weighed and mixed with 150 µl of ion chromatography water of the chosen pH. In contrast to the solution used in the heat flow cell experiments, the pH was not re-adjusted, exploring the amount of phosphate (and other salts) leached upon incubation at a given initial pH. After initial vortexing, leaching took place at a controlled temperature (T100 Thermal Cycler, Bio-Rad Laboratories). After the experimental incubation time, the samples were cooled to room temperature, vortexed, and centrifuged. The particle-free supernatant was then diluted with IC grade water for IC measurement.

