## Supplementary Information for "Heat flows solubilize apatite to boost phosphate availability for prebiotic chemistry"

### *Supplementary Figures*

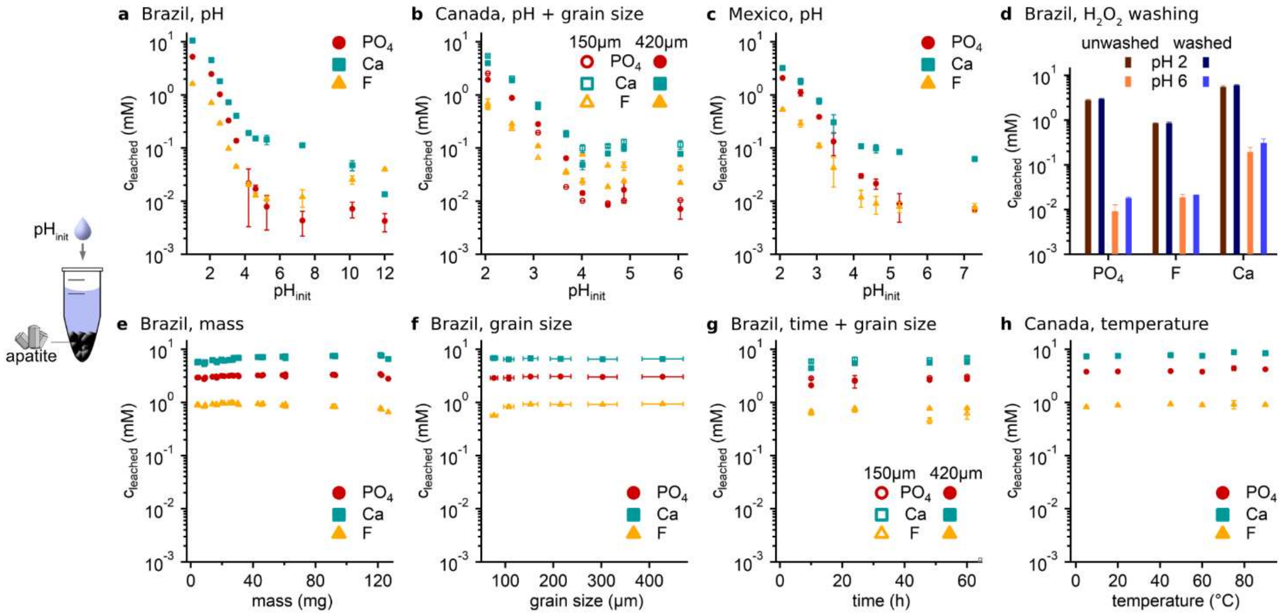

***Supplementary Figure 1.*** *Characterization of leaching of natural apatites.*

*Leaching was done for 60 h (if not stated otherwise) at 60 °C (if not stated otherwise) and for 30 mg of grains of diameter 355-500 µm in 150 µl of ion chromatography water. After the experimental time, ion concentrations were analyzed using ion chromatography (see Methods). All experiments were done in triplicate with error bars indicating the standard deviation.* ***(a-c)*** *Different natural apatites (for compositions, see Supplementary Table 1) were leached at various initial pH values. Concentrations and their ratios show variation over pH but remain similar for the three different apatites.* ***(d)*** *Washing of grains with $H_2O_2$ does not change the composition and concentration of leachates either.* ***(e-h)*** *Neither mass-to-volume ratio nor temperature, grain size, or time changes leached concentrations and ratios between species.*

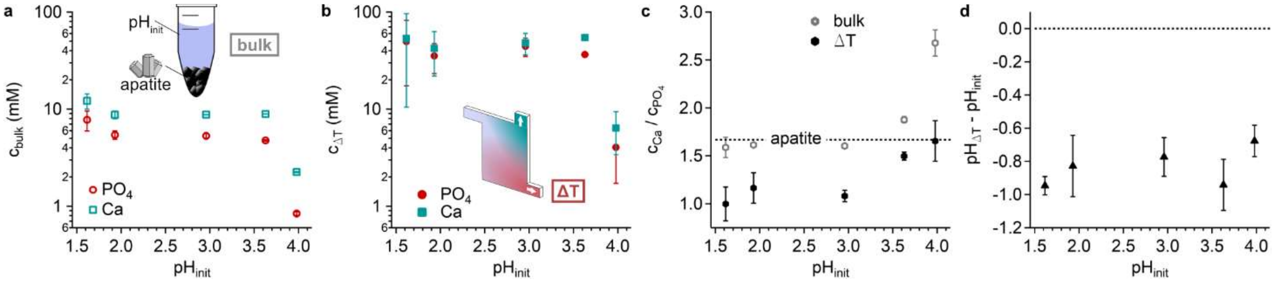

***Supplementary Figure 2.*** *Differential accumulation of calcium and phosphate from acidic-dissolved apatite.*

***(a)*** *Apatite samples (Brazil, see Supplementary Table 1) were acidic-dissolved by repeated addition of HCl (see Methods). When equilibrium was reached, and the pH stayed constant, samples were diluted with three fractions of water, previously adjusted to the same pH. Then, ion composition was measured using IC, showing the pH-dependent dissolution profile.* ***(b)*** *Flow through a microfluidic chamber that was exposed to a temperature gradient accumulates both phosphate and calcium but with different strengths.* ***(c)*** *Thereby, the initial 5:3 Ca:$PO_4$ ratio is altered to 1:1.* ***(d)*** *This ionic change is balanced by a shift in pH, thus keeping local charge neutrality.*

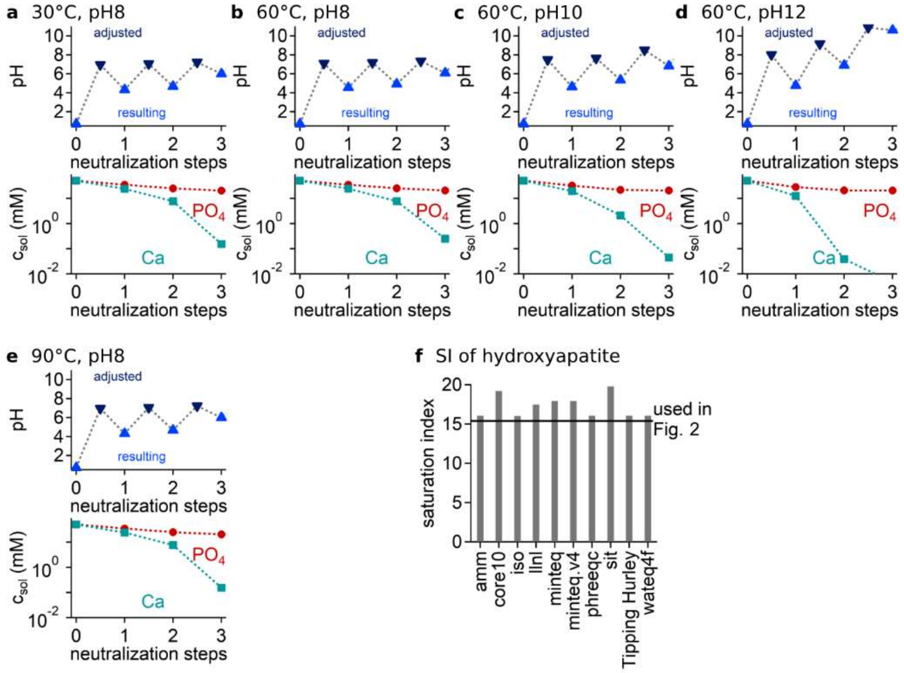

***Supplementary Figure 3.*** *Precipitation under different temperatures and pH.*

***(a-e)*** *In the experiments, we incubated samples for precipitation at 60 °C and set pH 8 in each step. Different temperatures and pH values did not change the dynamics; only the final pH values were modified.* ***(f)*** *Starting from a composition (50 mM P as $PO_4$, 50 mM Ca, 20 mM Cl (measured via IC, from acidic dissolution) and addition of 300 mM NaOH, see Methods) as in Fig. 1, we compared different databases for the saturation index ($SI = log(IAP/Ksp)$ with IAP the ion activity product and $K_{sp}$ the solubility product) of hydroxyapatite. Results were very similar and validated our method, which shows slightly inferior saturation indices.*

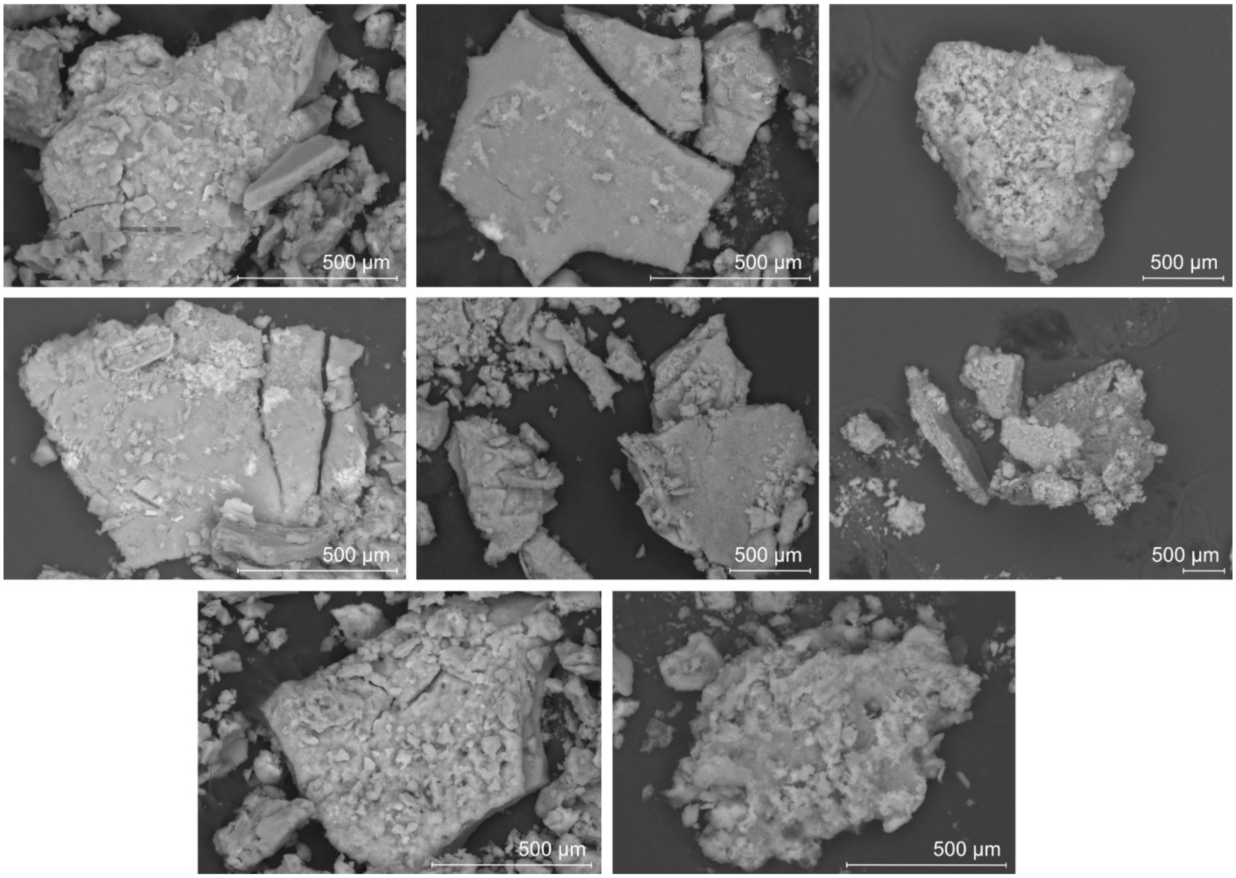

***Supplementary Figure 4.*** *Precipitates after re-neutralization.*

*Precipitation of solutions under SEM (see Methods) upon neutralization, CaO:$P_2O_5$ ratios are given in Supplementary Table 2.*

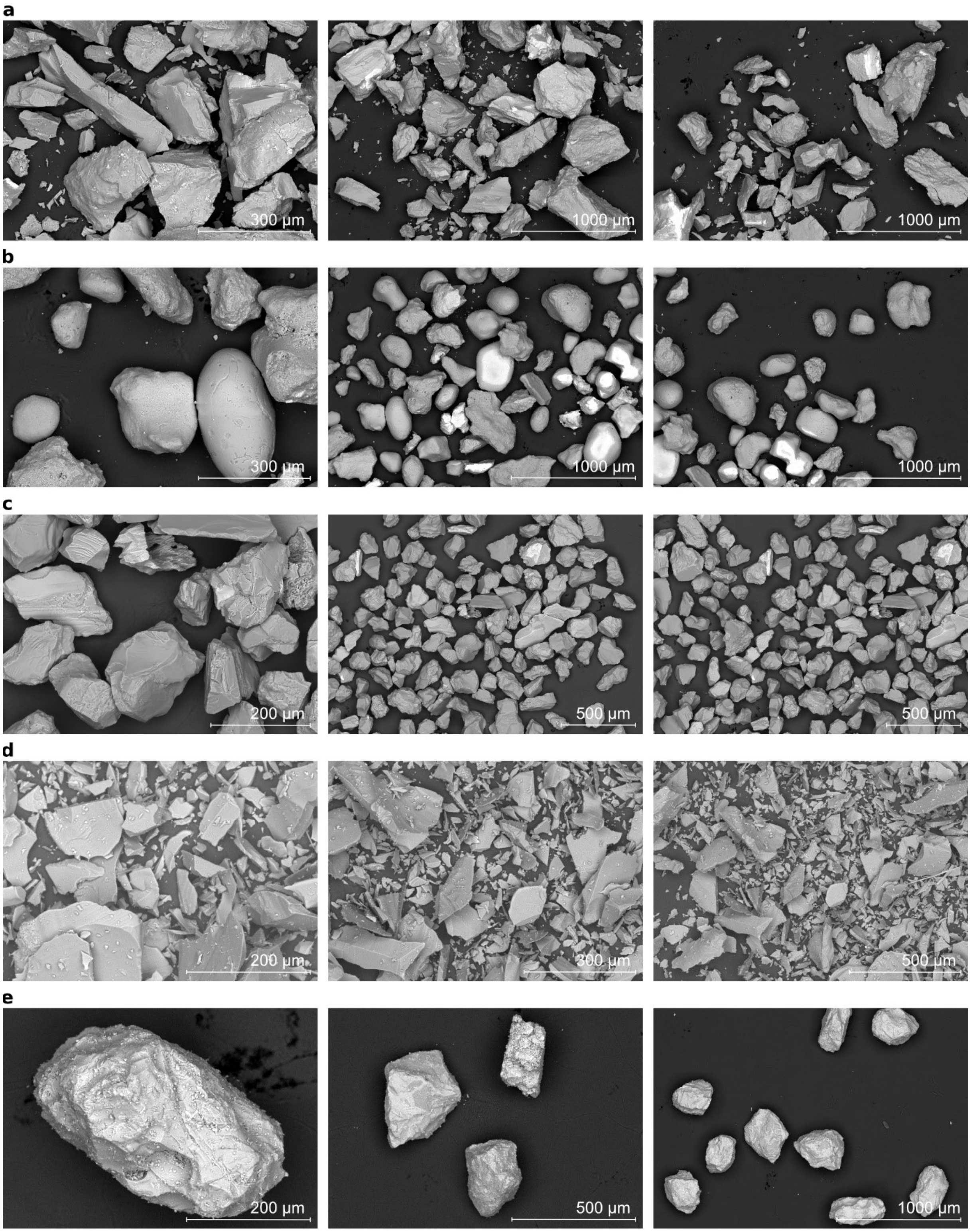

***Supplementary Figure 5.*** *SEM images of geomaterials tested for phosphate leaching and phosphate polymerization Part 1.*

*Compositions are given in Supplementary Table 3.* ***(a)*** *BSS: Basalt sand,* ***(b)*** *CAS: Carbonate sand,* ***(c)*** *SCS: Siliclastic sand,* ***(d)*** *VCG: Volcanic glass,* ***(e)*** *BF2: Basalt F2*

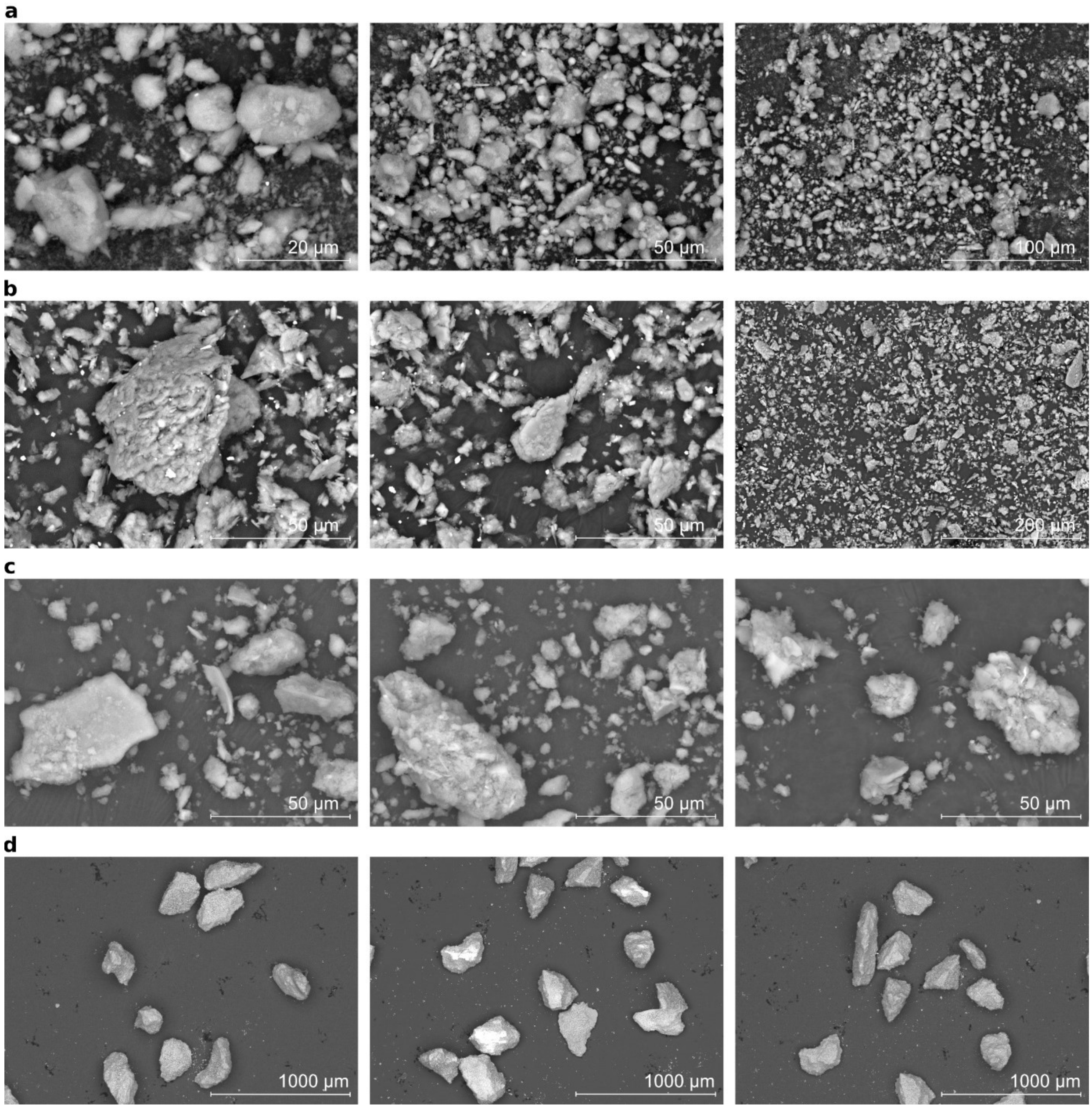

***Supplementary Figure 6.*** *SEM images of geomaterials tested for phosphate leaching and phosphate polymerization Part 2.*

*Compositions are given in Supplementary Table 3.* ***(a)*** *ILL: Illite,* ***(b)*** *KAO: Kaolinite,* ***(c)*** *MON: Montmorillonite,* ***(d)*** *ZEO: Zeolite*

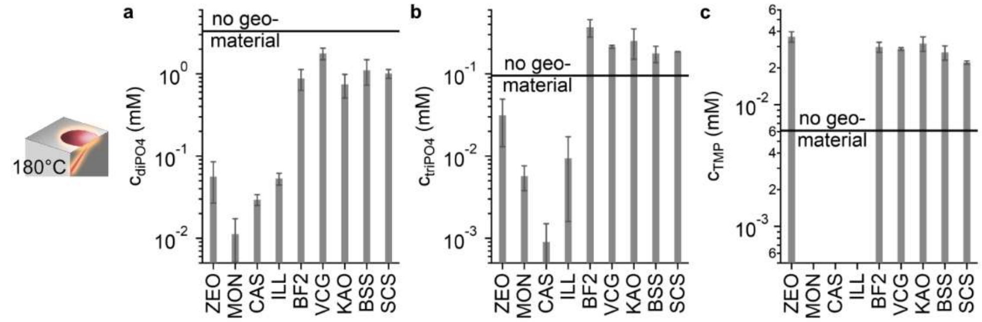

***Supplementary Figure 7.*** *Polymerization of phosphate on geomaterial.*

*10µl of 10 mM $NaH_2PO_4$ adjusted to pH 7 were added to 30 mg of geomaterial in glass vials and heated at 180 °C for three days. Analysis of redissolved phosphate species* ***(a-c)*** *shows clear positive or negative influence of geomaterial surface, which proved to be especially beneficial for the formation of trimetaphosphate (TMP,* ***c****). ZEO: Zeolite, MON: Montmorillonite, CAS: Carbonate sand, ILL: Illite, BF2: Basalt F2, VCG: Volcanic glass, KAO: Kaolinite, BSS: Basalt sand, SCS: Siliclastic sand.*

***Supplementary Tables***

| Abbreviation | Origin | CaO (%) | $P_2O_5$ (%) | $SiO_2$ (%) | $Na_2O$ (%) | $Fe_2O_3$ (%) | F (ppm) | Cl (ppm) |
|---|---|---|---|---|---|---|---|---|
| Brazil | Ipirá complex | 54.63 | 39 | 1.45 | 0.03 | 0.01 | 2.88 | 0.15 |
| Canada | Ontario | 55.05 | 41.53 | 0.17 | 0.19 | 0.02 | 2.42 | 0 |
| Mexico | Durango | 54 | 41.36 | 0.27 | 0.29 | 0.04 | 3.21 | 0.3 |

***Supplementary Table 1.*** *Compositions of apatite samples from different locations. Measurement was done using XRF analysis as described in the Methods.*

| Name | # of point measurements | $CaO$ / $P_2O_5$ (average) | $CaO$ / $P_2O_5$ (stdev) |
|---|---|---|---|
| Apatite Brazil | 17 | 1.40 | 0.02 |
| Apatite Canada | 6 | 1.33 | 0.02 |
| Apatite Mexico | 10 | 1.31 | 0.02 |
| Experimental (in, initial pH 1.9) | 21 | 1.53 | 0.19 |
| Experimental (bottom, initial pH 1.9) | 7 | 1.38 | 0.16 |
| Experimental (in, initial pH 1.9) | 5 | 1.52 | 0.05 |
| Experimental (bottom, initial pH 1.9) | 13 | 1.39 | 0.29 |
| Experimental (in, initial pH 3.6) | 65 | 1.21 | 0.07 |
| Experimental (bottom, initial pH 3.6) | 64 | 1.39 | 0.15 |

***Supplementary Table 2.*** *$CaO/P_2O_5$ ratios of precipitates in re-neutralized samples. Measurements were performed as described in the Methods. We assume errors increased due to small amounts of sample material and the presence of hydroxides, as shown before* [1,2].

| #M | Mate-rial | Na (%) | Mg (%) | Al (%) | Si (%) | P (%) | S (%) | K (%) | Ca (%) | Ti (%) | Mn (%) | Fe (%) |
|---|---|---|---|---|---|---|---|---|---|---|---|---|
| 24 | SCS | 2.45 | 0.06 | 10.5 | 79.2 | 0.23 | 0.02 | 2.52 | 1.61 | 0.10 | 0.03 | 2.98 |
| 25 | ILL | 0.16 | 3.56 | 23.3 | 50.1 | 0.37 | 0.04 | 5.52 | 8.71 | 0.72 | 0.10 | 6.84 |
| 29 | CAS | 0.65 | 1.79 | 0.37 | 0.00 | 0.05 | 0.70 | 0.05 | 96.2 | 0.03 | 0.02 | 0.05 |
| 32 | KAO | 0.03 | 0.15 | 42.7 | 54.4 | 0.44 | 0.07 | 0.82 | 0.09 | 0.29 | 0.02 | 0.51 |
| 30 | BSS | 4.07 | 2.43 | 19.7 | 60.8 | 0.07 | 0.04 | 1.96 | 5.23 | 0.33 | 0.10 | 4.80 |
| 27 | VCG | 3.61 | 3.51 | 14.4 | 49.8 | 0.49 | 0.08 | 0.97 | 8.92 | 4.17 | 0.19 | 13.6 |
| 23 | MON | 0.21 | 1.35 | 15.7 | 77.3 | 0.03 | 0.03 | 2.53 | 0.18 | 0.33 | 0.02 | 1.87 |
| 32 | ZEO | 0.24 | 0.89 | 12.9 | 77.2 | 0.02 | 0.03 | 3.49 | 2.83 | 0.22 | 0.01 | 1.83 |

***Supplementary Table 3.*** *Composition of geomaterials tested for phosphate leaching and phosphate polymerization.*

*Measurements were performed as described in the Methods using EDX, the corresponding SEM images are shown in Figs. S4-S5. #M describes the number of point measurements. SCS: Siliclastic sand, ILL: Illite, CAS: Carbonate sand, KAO: Kaolinite, BSS: Basalt sand, VCG: Volcanic glass, MON: Montmorillonite, ZEO: Zeolite.*

***Supplementary references***

1. Kleine-Boymann, M. *et al.* Discrimination between biologically relevant calcium phosphate phases by surface-analytical techniques. *Appl. Surf. Sci.* **309**, 27–32 (2014).
2. Miculescu, F. *et al.* Considerations and Influencing Parameters in EDS Microanalysis of Biogenic Hydroxyapatite. *J. Funct. Biomater.* **11**, 82 (2020).
